# Base pair editing of goat embryos: nonsense codon introgression into *FGF5* to improve cashmere yield

**DOI:** 10.1101/348441

**Authors:** Guanwei Li, Shiwei Zhou, Chao Li, Bei Cai, Honghao Yu, Baohua Ma, Yu Huang, Yige Ding, Yao Liu, Qiang Ding, Chong He, Jiankui Zhou, Ying Wang, Guangxian Zhou, Yan Li, Yuan Yan, Jinlian Hua, Bjoern Petersen, Yu Jiang, Tad Sonstegard, Xingxu Huang, Yulin Chen, Xiaolong Wang

## Abstract

The ability to alter single bases without DNA double strand breaks provides a potential solution for multiplex editing of livestock genomes for quantitative traits. Here, we report using a single base editing system, Base Editor 3 (BE3), to induce nonsense codons (C-to-T transitions) at four target sites in caprine *FGF5.* All five progenies produced from microinjected single-cell embryos had alleles with a targeted nonsense mutation and yielded expected phenotypes. The effectiveness of BE3 to make single base changes varied considerably based on sgRNA design. Also, the rate of mosaicism differed between animals, target sites, and tissue type. PCR amplicon and whole genome resequencing analyses for off-target changes caused by BE3 were low at a genome-wide scale. This study provides first evidence of base editing in livestock, thus presenting a potentially better method to introgress complex human disease alleles into large animal models and provide genetic improvement of complex health and production traits in a single generation.

## Introduction

Single nucleotide polymorphisms (SNPs) are the most common type of causative polymorphism in human and animal genomes for many genetic diseases and phenotypic changes to morphology. Efficient introduction of multiple causative SNPs in a single generation of livestock breeding to either introduce variants for complex genetic diseases or cause significant deviation from the phenotypic mean of a multigenic production trait holds substantial promise to developing better human disease models [1], or to provide rapid genetic improvement [2], respectively. To date, all previous reports on use of site-directed nucleases in livestock to achieve singlebase replacement have relied on double strand breaks (DSBs) and the stereotypical inefficiencies of the homology directed repair (HDR) processes. Even for monogenic traits, these low HDR efficiencies can be cost prohibitive for commercial use of gene editing in elite food animal populations, when combined with the expense of producing live animals from IVF embryos or somatic cell nuclear transfer [3].

Recent advances in genome editing using the type II bacterial clustered, regularly interspaced, palindromic repeats (CRISPR)-associated (Cas) system have enabled efficient modification of the genomes of many organisms, including livestock used for biomedical models or food [4–6]. However, in the latter case, the potential for unintended off-target mutations caused by site-directed nucleases remains an overemphasized concern for regulatory approval of gene edited animals and their offspring as food, even though the rate of natural mutagenesis in non-edited, bovine embryos can be increased three to four-fold during in vitro manipulation and maturation [7]. If the inefficiencies of normal IVF methods, editing by HDR, and additional physiological limits of total donor template concentration on embryo viability are also considered; then the promise of a base editing (BE) system to overcome the challenges of editing for multigenic changes in a single generation becomes quite promising. Recently, a simple BE system, which induces C to T conversion without any DSBs was described [8], and reports of successful application in human embryos [9–11], mice [12] and crops [13–16] have been documented. However, reports of using a BE system to introgress causative variation into livestock clones or embryos are still lacking.

Because the base editing efficiency of BE3 (rAPOBEC1-nCas9-UGI) was shown to be better than BE1 (rAPOBEC1-dCas9) and BE2 (rAPOBEC1-dCas9-UGI) [12], we chose BE3 to induce nonsense mutations into the coding sequence of our caprine target gene, *FGF5*. The encoded protein of *FGF5* is secreted during the hair growth cycle to signal inhibition of hair growth by a mechanism that blocks dermal papilla cell activation [17]. *FGF5* is regarded as the causative gene responsible for the *angora* phenotype (long hair) in mice [18], and we have previously shown that disruption of *FGF5* via CRISPR/Cas9 resulted in longer hair fibers and a 30% increase in cashmere yield per animal [19,20]. In the present study, we demonstrate that the BE3 system can achieve high efficiency single base substitution in *FGF5*, when introduced by microinjection into single cell embryos. We further examined the phenotypes at different morphological levels, and expected phenotypes were obtained even with the presence of mosaic expression of *FGF5* genotypes. By comparing the sequence of five mutant animals and four controls, we highlight that the BE3 induced off-target mutations are rare. Our work sets a foundation for improving the base editing approach for multigenic modification of microinjected embryos to produce better large animal models for complex human disease and provide a means to increase genetic gain for multigenic production and health in food animals.

## Results

### BE3 system induces base conversion in goat

BE3 [12] was used to introgress C to T transitions that produced non-sense codons in the coding sequence of caprine *FGF5*. To achieve this, the first exon of *FGF5* was scanned for potential target sequences, and four sites (three glutamine codons and one tryptophan codon) within exon 1 were identified as targets for a single C-to-T transition that would produce nonsense codons (Fig. 1a; Table S1).

**Fig. 1.**
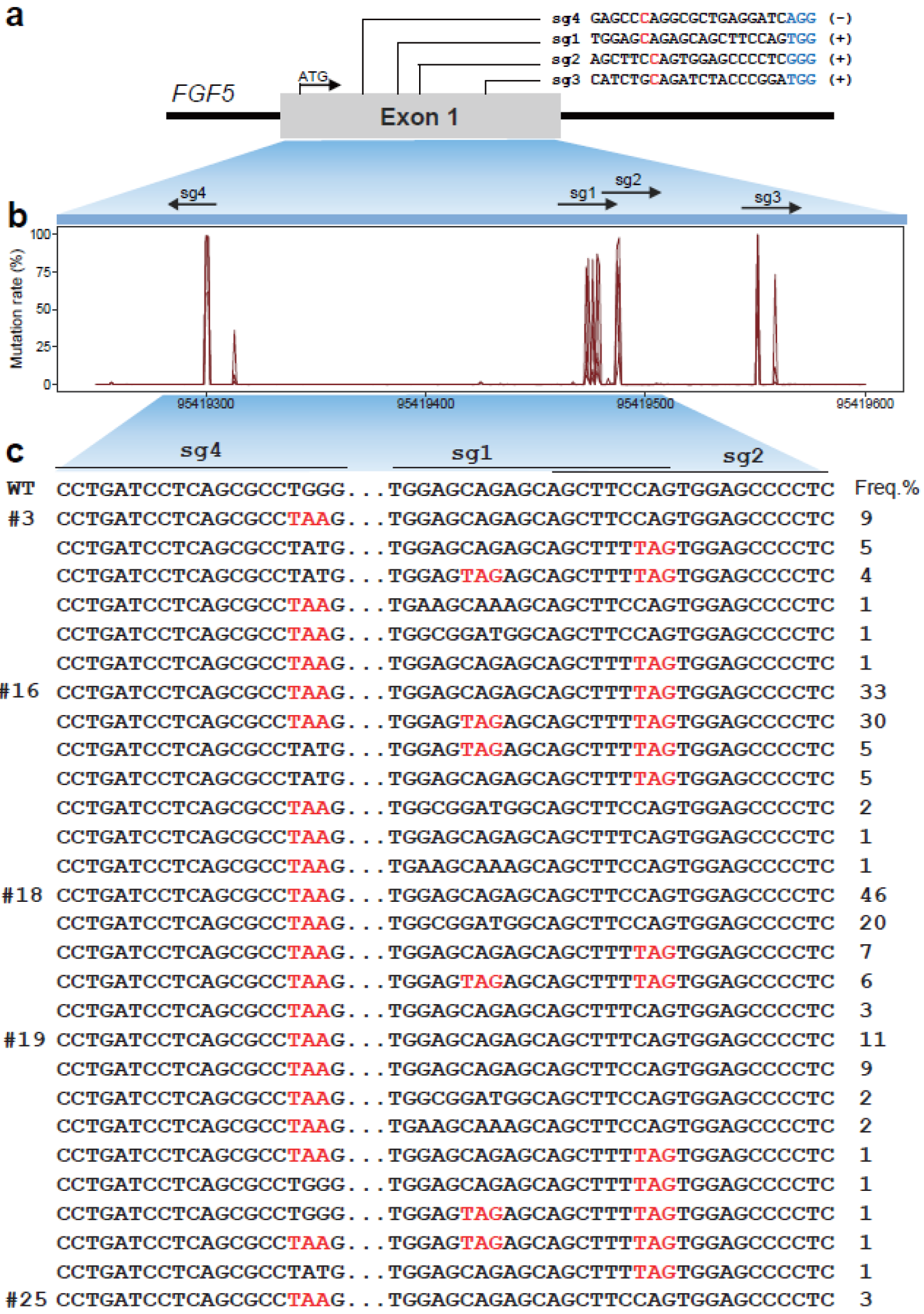
Generation of goats with cytidine-deaminase-mediated base editing. **(a)** Schematic diagram of target site at the *FGF5* locus. sgRNAs sequences are presented in red. PAM sequences are highlighted in blue and underlined. The BE3-mediated nucleotide substitutions are marked. **(b)** SNP rate at targeted regions in FGF5. **(c)** Alignment of sequences derived from deep sequencing in five edited animals (#3, #16, #18, #19, and #25). The target sequence is highlighted in red.

Initially, we transfected BE3 plasmid and single guide RNAs (sgRNAs) into goat fibroblasts to determine conversion efficiencies of the four sgRNAs before deploying into embryos. Extensive screening revealed none of the sgRNAs were effective at BE3 induced targeted base changes in *FGF5* (data not shown). Even with this negative result, we moved ahead to test BE3 efficiency in injected single cell embryos. A total of 48 single cell zygotes were surgically collected from five naturally mated donor ewes. After micro-injection of BE3 mRNA and sgRNAs mixtures, 22 surviving embryos (two-cell stage) were transferred to seven surrogate mothers. A total of five lambs (10% of total embryos) from three surrogate females were successfully delivered (Table 1). Based on Sanger sequencing of exon 1 from these five animals, each animal had accumulated alleles of *FGF5* with at least one nonsense mutation or other mutational types induced by the BE3 system (Figure S1).

**Table 1.**
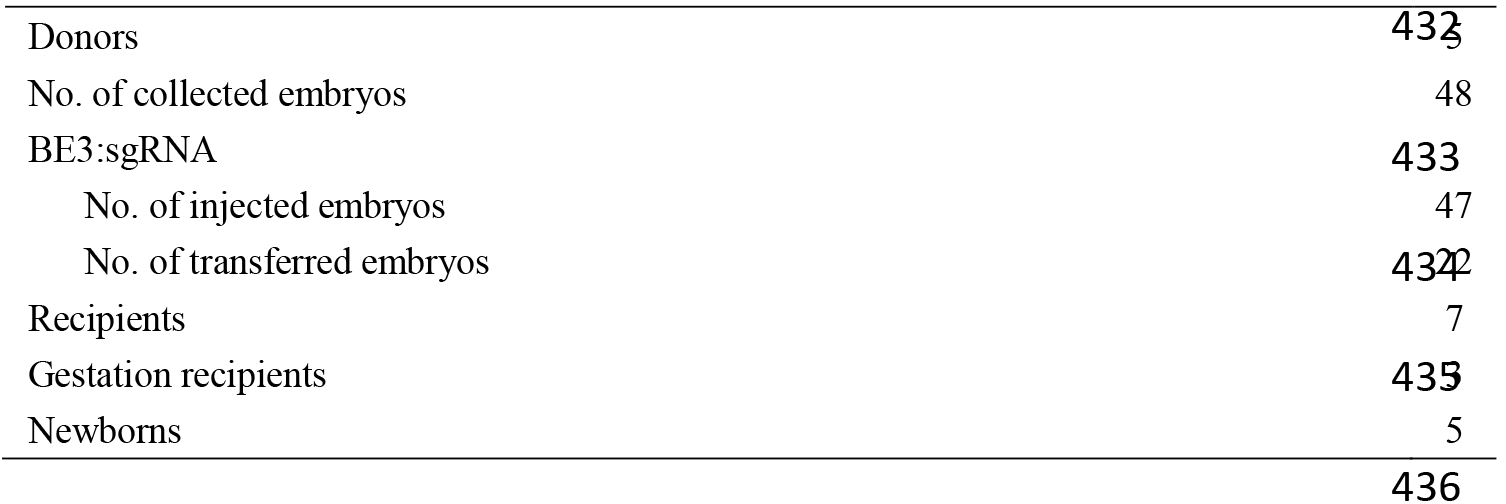
Summary of the goats generated with BE3-mediated base editing.

To fully characterize the genotypes/haplotypes at the target sites, genomic DNA samples were subjected to deep sequencing of *FGF5* exon 1, which allowed determination of cumulative and individual BE3 conversion efficiencies for each sgRNA. The targeting efficiency between the four target sites (sg1, sg2, sg3, and sg4) varied considerably (Fig. 1b). Overall, 13 different base substitution mutations were observed at or nearby the four targeted sites, including C-to-A and C-to-G conversions, and all five founder animals were mosaic (Fig. 1c; Figure S2). The highest frequency of introgressed nonsense alleles was found in animals 16 and 18 (both >75%), while animals 25 had cumulative nonsense allele frequencies lower of 3% (Fig. 1c). Sg1 was the only guide RNA to mutate alleles with the intended single base change; albeit at a low frequency across all sequence reads (~9%). Sg2 had a cumulative conversion rate of 20%, but this change had an additional mutation at upstream flanking base (C to T transition), which resulted in a silent codon mutation. The sg4 guide RNA had the best rate of conversion to nonsense alleles (39% across all reads). Interestingly, this was the only sgRNA designed against the non-transcribed strand of FGF5. Additionally, a second conversion event was always present at a flanking base pair, similar to sg2. Thus, the non-sense conversion was to an unintended TAA nonsense codon instead of the designed TAG codon. For all the sgRNAs, unintended mutations were commonly found. At the sg3 site, the targeted C-to-T transition was not observed, however, a Leu102Met missense mutation 3 bp upstream of the target site was found at 33% allele frequency across all sequence reads (Figure S2). In addition, we also observed a very low frequency of small indels flanking the sgRNA binding sites (Fig. 2a).

**Fig. 2.**
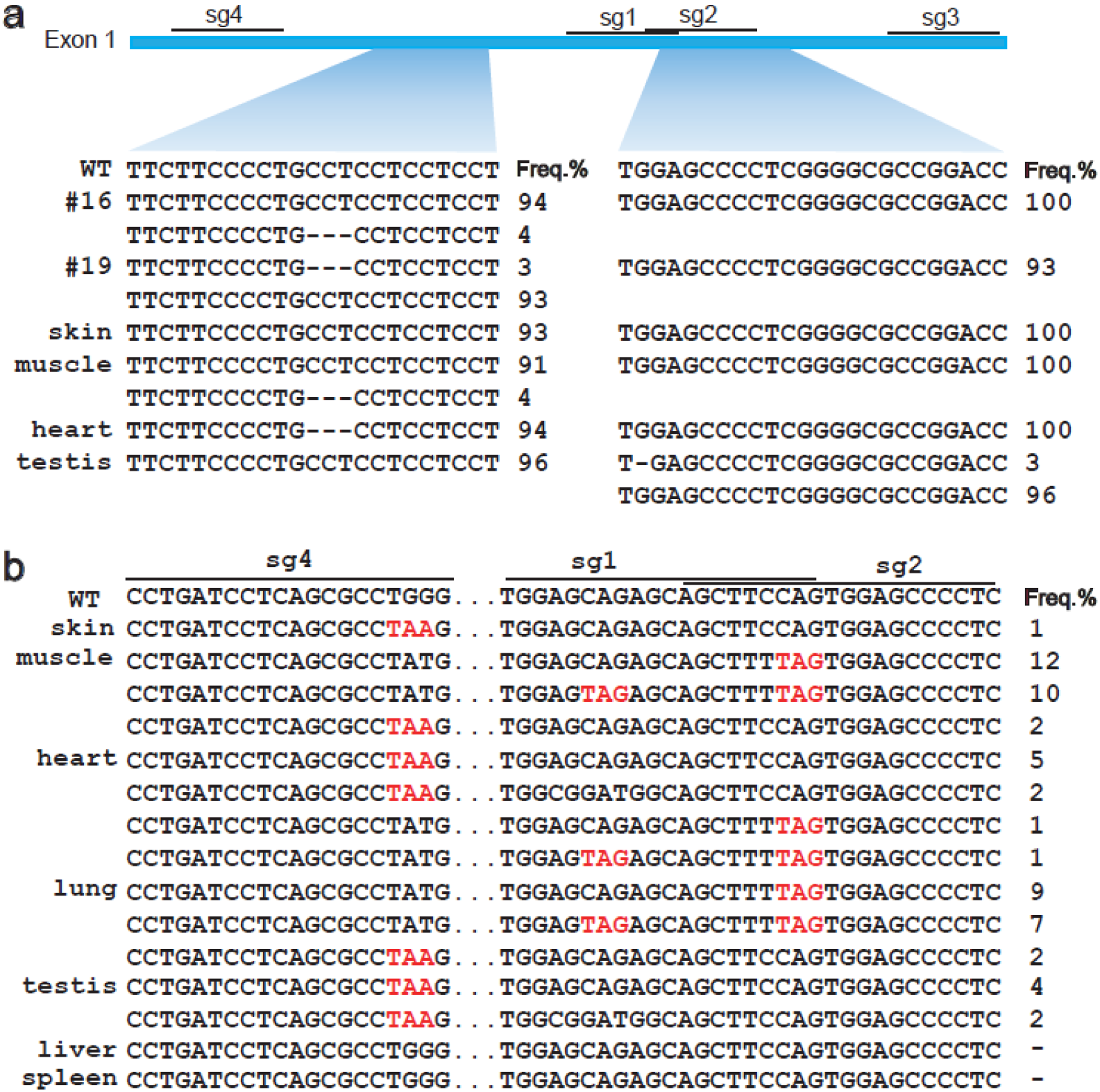
indel distribution and mosaic analyses of mutant sequences. **(a)** indels identified within the sequenced regions in four mutant animals (#16, #18, #19, and #25) or in tissues of #3. **(b)** Mosaic analyses of mutant sequences at four target sites in seven different tissues (skin, muscle, heart, lung, testis, liver, and spleen) of founder animal #3. No stop codon mutations were found in liver and spleen samples.

To examine the extent of mosaic modification within an animal, we sequenced the four sgRNA sites of animal #3 using seven tissues (muscle, heart, liver, spleen, lung, skin, and testis) that represented all three germ layers. Approximately 10,000 randomly sampled sequences at each target site were clustered. Deep sequencing revealed that the mosaic ratios of mutant FGF5 alleles were relatively similar across different tissues (Fig. 2b; Figure S2), and nonsense mutations at the sg2 and sg4 sites were not found in sequences from liver and spleen tissues. Collectively, our results highlighted that some combinations of BE3 and sgRNA can reach a high efficiency of targeted conversion through direct microinjection of zygotes. However, we also observed lack of proper targeting and unintended allele complexity, which seemed to be dependent on sgRNA design.

### Functional validation of BE3-mediated base editing

Given that *FGF5* is known to inhibit hair growth by blocking dermal papilla cell activation [17], and is determinant of hair length in dogs [21], cats [22], mice [18], and humans [23]; then characterization of fibers in founding animals would reveal the penetrance of the mosaic genotypes on phenotypes. First, the length of hair fibers (outer coat hair and inner fine fiber) between mutant and control animals was measured and compared. The fibers from BE3 produced animals were significantly longer than the control animals (p <0.05, Student’s *t*-test) (Fig. 3a). If animal #25 was removed from this analysis due to its much lower allele frequency of nonsense codons, then the phenotypic differences between edited and control animals would have been even greater. Next, we completed a histological analysis comparing skin tissues from four mutated live animals (#16, #18, #19, and #25) and the corresponding wildtype animals. Hematoxylin and eosin (H&E) staining indicated that there were more secondary hair follicles (SHF) in the skin of BE3 edited goats (Fig. 3b; Figure S3). Considering our previous report [20], these results confirm that inducing point mutations through BE3 yielded a similar phenotype to site-directed nuclease gene disruption.

**Fig. 3.**
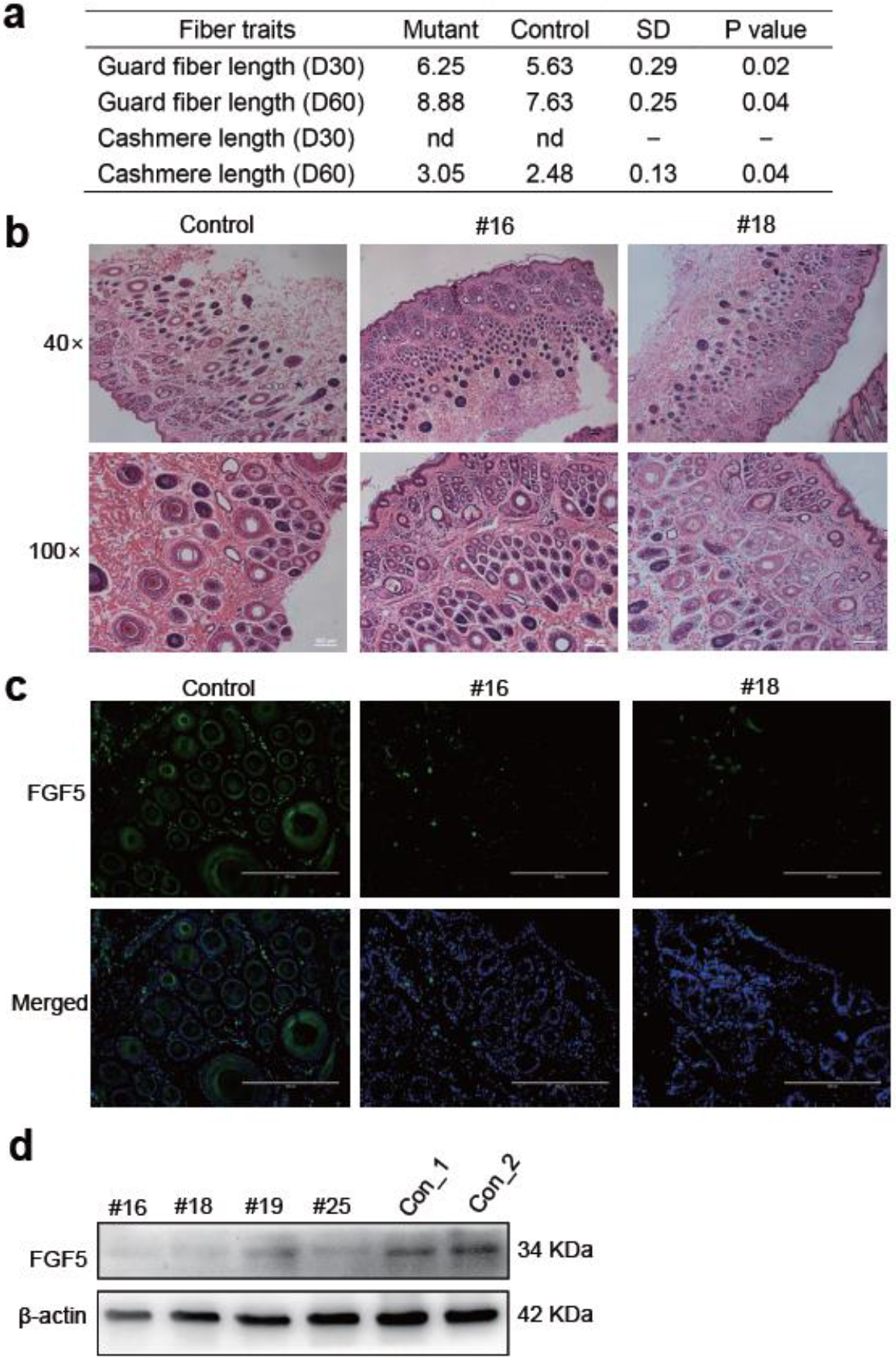
Phenotypic changes of mutant goats with BE3-mediated base editing. **(a)** Hair fiber phenotypes in mutant and wildtype. **(b)** H&E staining demonstrating hair follicle morphology in the skin tissues of mutant (#16 and #18) and wildtypes. **(c)** The skin tissues from mutant (#16 and #18) and wild-types were subject to immunofluorescence staining using anti-FGF5 antibodies (green) and counterstained with Hoechst33342 (blue). Scale bar, 200 μm. **(d)** Western blot analysis using anti-FGF5 (34 kDa) and anti-β-actin (control) antibodies.

Finally, we examined FGF5 protein expression by immunofluorescence staining of skin samples from four mutant animals and two controls. The immunofluorescence showed that FGF5 expression was significantly reduced in animals derived from BE3-treated embryos (Fig. 3c; Figure S4); however, the location of the FGF5 protein was not altered between animal types (Fig. 3c; Figure S4). The specificity of immunofluorescence staining for FGF5 in skins was further confirmed by western blotting (Fig. 3d). All combined, we concluded that the observed phenotypes were caused by the nonsense mutations in *FGF5*.

### Off-target validation in mutant animals

To assess potential off-target effects induced by BE3, we predicted off-target sites via Cas-OFFinder [24]. A total of 19 off-target sites were predicted with up to three mismatches for the four sgRNAs (Table S2). We conducted targeted deep sequencing of PCR amplicons to screen the mutations at these 19 sites for founder animals #16 and #25, and observed high SNP substitution rate at sg2_OT1 site only in founder #16 (Fig. 4).We further sequenced the sg2_OT1 site using genomic DNA from #16 and its parents, the sequencing results confirmed the existence of the TT genotype in #16 and is not inherited from their parents (Figure S5), indicating this might be an off-target mutation induced by BE3.

**Fig. 4.**
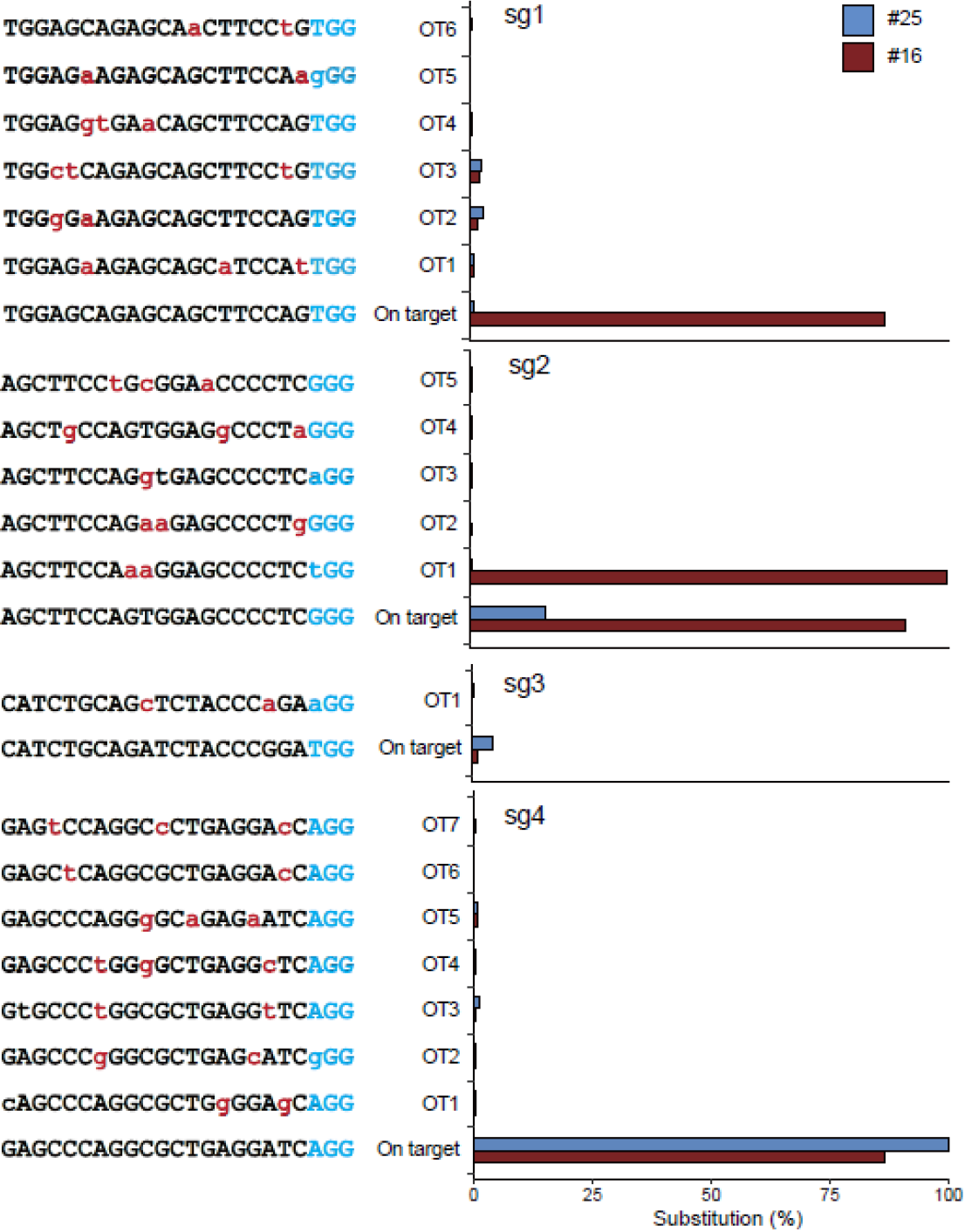
Off-target mutation detection of potential off-targeted sited predicted by Cas-OFFinder. Deep sequencing was used to measure substitution frequencies at predicted target sites for four sgRNAs in #16 and #25. Mismatched nucleotide and PAM sequences are indicated in red and in blue, respectively

To more extensively identify off-target sites throughout the goat genome, we conducted whole-genome sequencing (WGS) to identify BE3 off-target mutations in all five founder animals. We also sequenced four control animals (#1, #31, #61, and #92) (Table S3). After filtering all SNPs called by GATK and Samtools to subtract naturally-occurring variants in our goat SNP database (234 individuals from 11 breeds, ~79 million SNPs, unpublished data) and filtering SNPs found in the four control animals, an average of about 300,000 SNPs remained for each founder animal (Table 2; Table S3). Base substitutions other than C to T or A or G conversions were further excluded based on previously reported methods [12]. Next, we examined potential off-targets for ~200,000 predicted protospacer adjacent motif (PAM), which were predicted based on allowing five mismatches with each sgRNA. 20 sites, including the off-target mutation identified by Cas-OFFinder (Figure S5) in #16, were determined to contain SNP variations induced by BE3 in the four target sites at five mutant animals (Table S4), ~1 site was found at a single site for each animal. Of the 20 potential off-target sites, seven variants were not genetically inherited and were determined as unwanted off-target mutations (Figure S5&S6), representing a slight off-target mutagenesis in the edited animals with base editing.

**Table 2.**
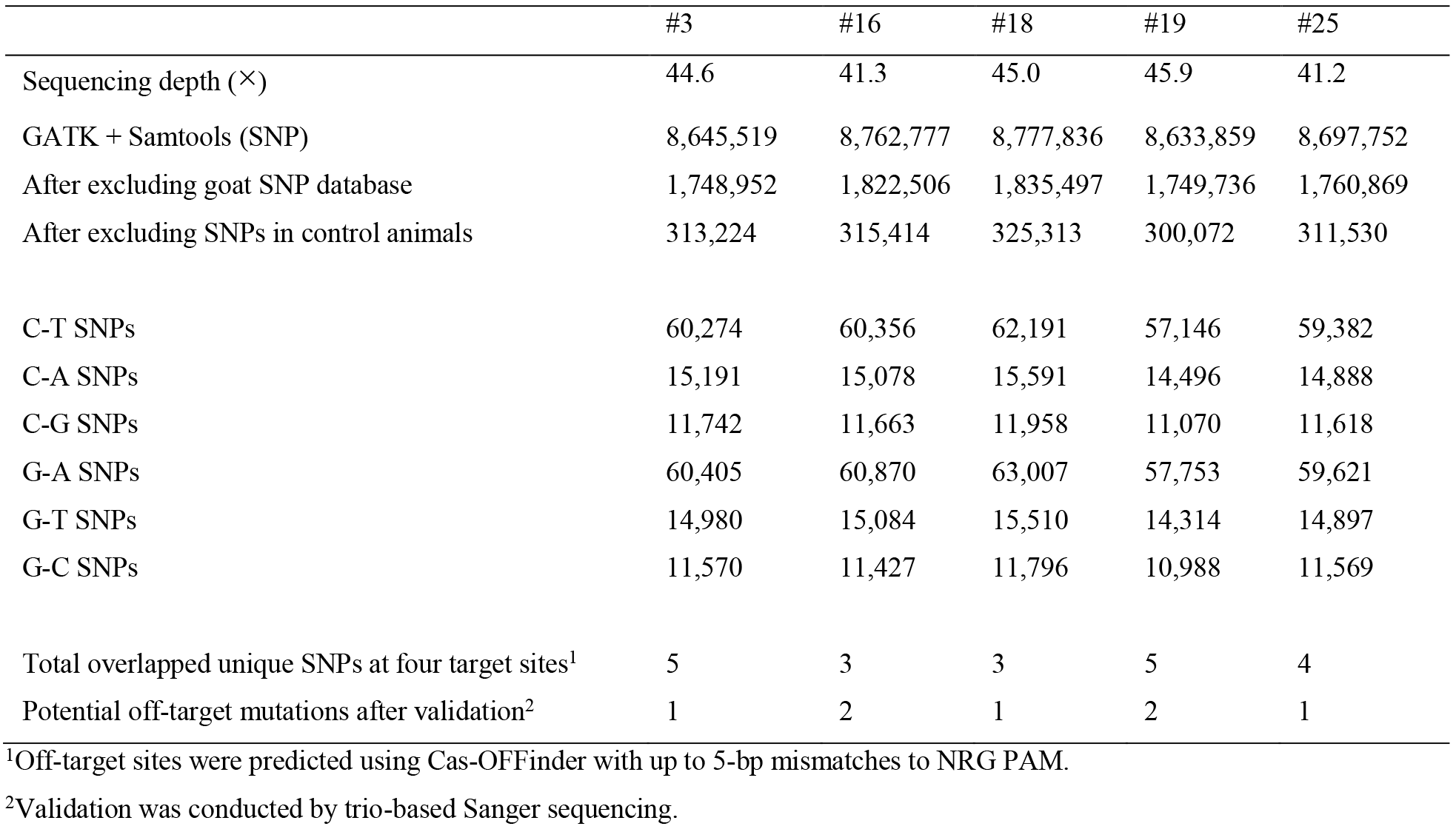
WGS analyses of mutant animals for revealing genome-wide off-target mutations.

## Discussion

The ability to introduce causative variant SNPs into naïve livestock populations for the purpose of genetic gain holds great promise for mitigating some of the challenges related to global food security. Numerous reports have demonstrated that single-base pair alterations can be directly engineered into the donor livestock genomes after a targeted DSB event with the aid of either a plasmid template or single-stranded oligodeoxynucleotides (ssODNs) to direct HDR processes. However, efficient nucleotide substitutions require reduced levels of non-homologous end-joining (NHEJ) to avoid unwanted indels. Thus, targeting efficiencies of single SNP substitutions by HDR remain relatively low [25,26]. These inefficiencies are both a practical and economic limitation to deployment of new traits in livestock, because advanced reproductive techniques are already inefficient and expensive, sourcing of elite genetics for editing is also costly, and generation intervals to test outcomes and allele transmission are much longer than those for rodent models.

The recently emerged RNA-guided programmable deaminase [8], provides another potential tool for genetic improvement using new breeding technologies. Specifically, the possibility to convert specific bases without generating DSBs and adding additional nucleic acid material to guide HDR makes multigenic editing a more tangible possibility for livestock. Here we report for the first time the use of a base editor in livestock embryos to induce nonsense codons in a gene (*FGF5*), known to have observable and definitive effects on hair fibers in mammals. However, since we employed four sgRNAs within a short genomic region, it is possible there was some competitive interference for base editing between sg1 and sg2. Moreover, we found very little off-target effects, which provides strong support for the reliability of base editor techniques to produce large animals intended for biomedical studies and food production. Clearly, pre-optimized sgRNAs need validation for their base editing precision prior to deployment in cloneable cell lines and embryos.

The targeted site-specific mutagenesis with programmable nucleases (e.g. Cas9 and Cpf1) rely on zygote microinjection often results in mosaicism with respect to mutated cells [27–29]. Although Kim reported homozygous mice were obtained with microinjection of BE3 mRNA and sgRNA [12], we observed mosaicism in the founder animal #3 at all sgRNA sites by sequencing somatic tissues and testis representing three germ layers (Fig. 1c, Figure S4). The frequent mosaic patterns observed in our study might be caused by extended BE activity in the rapidly developing embryo and/or by the poor spreading of the introduced mRNA after zygote injection at 1-cell-stage, which likely have resulted in asymmetric mRNA accumulation resulting in mosaicism, as previously observed with ZFN, TALEN and CRISPR-mediated targeting [27, 28, 30]. Furthermore, the germline mutations observed in the testis of animal #3 indicated the edited variants will be transmitted to next generation; albeit in the case of this animal #3 at a low frequency (~20%).

To further investigate the unintended off-target mutation that may be produced by BE3 modification. We first screened predicted putative off-targets *in silico*, and one off-target mutation was revealed through deep sequencing (Fig. 4; Figure S5). To fully characterize the possible BE3 induced off-targets at genome-wide scale, we sequenced five mutant animals with a high coverage (>40 ×). Sequence comparison revealed a total of seven potential off-target mutations which were not inherited from their parents in five founders at four target sites (Figure S6), indicating these unwanted SNPs might be induced by BE3-mediated base editing. Along with previous studies in human embryos [9–11] and mice [12], our results indicated a low incidence of BE3-induced off-target mutagenesis, and the off-target mutations are depending largely on sgRNA design. Therefore, it is highly recommended to pre-screen the efficient functional sgRNAs without potential off-target mutations.

In this study, we succeeded in generating single-base pair substitutions using zygote injection of BE3 modification in goats with reasonable efficiency depending on sgRNA design (up to 39% for sg4) and low indel rates. However, a high mosaicism upon mutation induction was observed. With the rapid development of base editor tools, which are able to mediate almost all base type substitutions [8, 31], and to eliminate mosaicism derived by zygote injection [32], the base editor mediated genome editing will greatly accelerate the gene correction and validation/elucidation of functional SNPs.

Taken together, we demonstrated the BE3-mediated base editing of four target sites in goats. We further examined the phenotypic and genetic changes to investigate the consequence of base pair editing, and provided strong support that the BE3 induced off-target mutations are rare at genome-wide level. Our attempts in goats opens up unlimited possibilities of genome engineering in large animals for applications in agriculture and biomedicine.

## Methods

### Animals

Five rams (2–3 years old, body weight: 30–50 kg) and 12 ewes (5 donors and 7 recipients, 2–3 years old, body weight: 24–40 kg) were used in the present study. The animals were regularly maintained in the Shaanbei Cashmere Goat Farm of Yulin University. All the protocols involving animals were approved by the College of Animal Science and Technology, Northwest A&F University (Approval ID: 2016ZX08008002).

### mRNA and sgRNA preparation

BE3 was reported previously [8] and was obtained from Addgene (plasmids 73021). *In vitro* transcription of BE3 and sgRNAs were conducted with some modifications. Briefly, the BE3 plasmid was extracted with plasmid midi kit (TIANGEN, DP107-02), and linearized by digestion with Bbs I (NEB, R0539S). The linearized plasmid was then purified with PCR purification kit (Axygen, AP-PCR-500G) and in vitro transcribed using mMESSAGE mMACHINE T7 Ultra Kit (Ambion, Life Technologies, AM1345). sgRNAs (Table S1) were amplified from the constructed Puc57-T7 sgRNA plasmid (Addgene plasmids 51132) with primers (F: 5’-TCTCGCGCGTTTCGGTGATGACGG-3’; R: 5’-AAAAAAAGCACCGACTCGGTGCCACTTTTTC-3’). The purified PCR products were then used as templates for transcription using the MEGAshortscript T7 Transcription Kit (Ambion, Life Technologies, AM1354). mRNAs and gRNAs were subsequently purified with the MEGAclear kit (Ambion, Life Technologies, AM1908).

### BE3/sgRNA efficacy test in goat fibroblasts

The fibroblasts were cultured for five passages in DMEM medium (Gibco) supplemented with 10% FBS (Gibco) and 1% penicillin-streptomycin (Gibco) until 80%–90% confluency, which were then subjected for transfection. The transfection procedures were conducted with Lipofectamine 3000 Reagent (Invitrogen) according to the manufacturer’s instructions. Briefly, fibroblasts were separately transfected with each sgRNA (2.5 μg for each sgRNA) along with 2.5 μg of BE3 plasmid by Lipofectamine 3000 in a 6-well culture plate for 48 h. 1.0 mg/mL puromycin was added to the medium (1:1000 dilution) and incubated for 72 h. Genomic DNA was isolated from fibroblasts for Sanger sequencing. Targeted fragments were amplified with 2xEasyTaq SuperMix (TransGen Biotech), then purified with a PCR cleanup kit (Axygen, AP-PCR-50).

### Generating of the edited animals

Healthy ewes with regular estrus cycles were chosen as donors for zygote collection. Zygotes were collected through surgical oviduct flushing from the donors by estrus synchronization and superovulation treatment as we described previously [19]. The mixture of BE3 mRNA (25 ng/μL) and sgRNAs (10 ng/μL for each sgRNA) was injected into surgically collected zygotes at one-cell-stage (~14 h post-fertilization) using the FemtoJect system (Eppendorf). The parameters of injection pressure, injection time and compensatory pressure were 45 kpa, 0.1 s and 7 kpa, respectively. Microinjection was conducted in manipulation medium TCM199 using the micromanipulation system ON3 (Olympus). The injected zygotes were transplanted into the ampullary-isthmic junction of the oviduct of the surrogate ewes after culturing in Quinn’s Advantage Cleavage Medium and Blastocyst Medium (Sage Biopharma) for more than ~24 h. Pregnancy was determined by observing the estrus behaviors of surrogates at every ovulation circle.

### WGS and data analysis

WGS was carried out using the Illumina Hiseq3000 at mean coverages of >40 × for five mutant goats (#3, #16, #18, #19, and #25) and 25 × for four control animals (#1, #31, #61, and #92). For each animal, genomic DNA was extracted from peripheral venous blood samples with a Qiagen DNeasy Blood and Tissue Kit (Qiagen). To construct the WGS library, 1 μg of genomic DNA was fragmented to around 300 bp by ultrasonication using a Covaris S2 system. Then, the sheared DNA fragments were used for library construction using an Illumina TrueSeq DNA library preparation kit at Novogene (www.novogene.com). The qualified reads were mapped to the goat reference genome (ARS1) [33] using BWA (v0.7.13) tools [34]. Local realignment and base quality re-calibration were conducted using the Genome Analysis Toolkit (GATK) [35]. SNPs and small insertion and deletions (indels) (< 50 bp) were called using both GATK [35] and Samtools [36].

The called SNPs from WGS were first filtered according to the following criteria: (1) SNPs that were identified by GATK and Samtools; (2) filtering SNPs that exist in a goat SNP database (n=234, 11 populations including 30 cashmere goats, > 79 million SNPs, unpublished data); (3) filtering common SNPs that were existed in the control groups. Of these remaining SNPs, we selected SNPs with C and G converted to other bases. The potential off-target sites were predicted with Cas-OFFinder [24], by considering up to five mismatches. We next compared the DNA sequences encompassing the SNP sites with the off-target sequences for each sgRNA.

### Targeted deep sequencing

Targeted genomic sites were amplified with the high-fidelity DNA polymerase PrimeSTAR HS (Takara) using primers flanking the sgRNA-target sites or predicted off-target sites (Table S5). The amplified PCR products were fragmented with a Covaris S220 ultrasonicator, and then amplified using the TruSeq CHIP Sample preparation kit (Illumina). After being quantified with a Qubit High-Sensitivity DNA kit, the PCR products with different tags were pooled and were conducted for deep sequencing with Illumina platform using standard protocols. Each sequenced site obtain > 3 M clean reads.

### Off-target mutation analysis

First, we screened for the off-target mutations by sequencing the predicted putative off-target sites derived from Cas-OFFinder analysis [24]. A total of 23 potential off-target sites were identified in the goat genome for the four sgRNA used in this study (Table S2), each of putative off-target had three nucleotide mismatches to the sgRNA target regions. PCR amplicons were carefully examined by Sanger sequencing and followed by targeted deep sequencing, primers were summarized in Table S5. Furthermore, WGS of five mutant animals (#3, #16, #18, #19, and #25) and four control animals (#1, #31, #61, and #92) were carried out to extensively examine off-target mutations at the genome scale.

### H&E and immunofluorescent staining

A portion of the skin tissues of mutant (#16, #18, #19, and #25) and WT animals were biopsied. The biopsies tissue was immediately fixed in 4% paraformaldehyde at 4 °C overnight. The fixed tissues were then embedded in paraffin using standard procedures for further H&E staining. The tissue sections were de-waxed, rehydrated, and stained using standard immunohistochemical protocols. The immunofluorescence staining was conducted with anti-FGF5 (Sigma-Aldrich, 1:300) primary antibody, the sections were then counterstained with Hoechst 33342 and analyzed by confocal laser microscopy.

### Western blot analysis

Skin samples were subjected to total protein extraction with a ProteoJET Membrane Protein Extraction Kit (Fermentas), and then quantified using the Bradford assay. Equal amounts of soluble protein were separated by SDS/PAGE and transferred onto a polyvinylidene difluoride membrane (PVDF, Roche). Immunoblotting was conducted using antibodies specific for FGF5 (Abcam, 1:1000) and anti-GAPDH (Abcam, 1:1000). Primary antibodies were visualized using a fluorescence imager system (Sagecreation). Variations in sample loading were corrected by normalizing.

### Data Availability

The raw data of sequenced animals involved in this study are available under BioProject ID: PRJNA470771 and SRA accession no. SRR6378096.

## Acknowledgments

We thank members of Chen lab for help with animal experiments. This work is supported by grants from National Natural Science Foundation of China (31772571, 31572369), the Key Research Program of Shaanxi Province (2017NY-072), the Tan Sheep Breeding Project from Ningxia (NXTS201601), as well as by China Agriculture Research System (CARS-39). XW was supported by the Innovative Talents Promotion Plan in Shaanxi Province, and the Tang Scholar Foundation.

## Authors’ contributions

BM, XH, YC, and XW conceived and designed the experiments. GL SZ BC HY JH YH YD YLiu YW YLi GZ and YY performed genotyping and phenotyping analyses. HY and QD injected embryos. JZ and YH prepared the mRNA and sgRNA. CL YJ CH and TS analyzed the sequencing data. XW BP and TS wrote the manuscript. All authors read and approved the final manuscript.

## Competing interests

The authors declare competing financial interests.

## Additional information

Supplementary Tables: Table S1-Table S5.

Supplementary Figures: Figure S1–Figure S3.

